# Spatio-temporal correlations between catastrophe events in a microtubule bundle: a computational study

**DOI:** 10.1101/694836

**Authors:** Makarand Diwe, Manoj Gopalakrishnan

## Abstract

We explore correlations between dynamics of different microtubules in a bundle, via numerical simulations, using a one-dimensional stochastic model of a microtubule. The GTP-bound tubulins undergo diffusion-limited binding to the tip. Random hydrolysis events take place along the filament, and converts GTP-tubulin to GDP-tubulin. The filament starts depolymerising when the monomer at the tip becomes GDP-bound; in this case, detachment of GDP-tubulin ensues and continues until either GTP-bound tubulin is exposed or complete depolymerisation is achieved. In the latter case, the filament is defined to have undergone a “catastrophe”. Our results show that, in general, the dynamics of growth and catastrophe in different filaments are coupled to each other; closer the filaments are, the stronger the coupling. In particular, all filaments grow slower, on average, when brought closer together. The reduction in growth velocity also leads to more frequent catastrophes. More dramatically, catastrophe events in the different filaments forming a bundle are found to be correlated; a catastrophe event in one filament is more likely to be followed by a similar event in the same filament. This propensity of bunching disappears when the filaments move farther apart.

## 1 Introduction

The dynamical instability of microtubule filaments, and in particular, the “catastrophe” transition, has been the subject of a large number of experimental, theoretical and computational studies[1–8]. Catastrophe refers to the sudden and abrupt transition of a growing microtubule to a shrinking state, which originates from the hydrolysis of guanine tri-phosphate (GTP) molecule bound to adsorbed tubulin. In vivo, the reverse “rescue” transition is also often observed, which resurrects a microtubule before it shrinks away completely. The combination of catastrophe and rescue transitions generates the characteristic zig-zag growth curves of microtubules. This unique dynamics of microtubules is crucial in facilitating the search and capture of chromosomes during cell division.

In many cells, a bundle of microtubules, rather than a single one, is found to attach to a single kinetochore. The number of filaments forming the bundle varies from one in budding yeast (*S. cerevisiae*) to 15-35 in mammals. In such situations, for efficient segregation of chromosomes, it is important that the the dynamics of different filaments are coordinated. Earlier experiments have indicated that a bundle of microtubules pushing against a common barrier undergoes collective catastrophes[9], which has also been studied in numerical simulations[10]. In this case, collective behaviour arises from the sharing of the total resistive force among all the filaments in the bundle.

But, is it possible for different microtubules in a bundle to interact with each other even in the absence of external force ? Experimental observations in *C. elegans* embryos [11] during metaphase show that astral microtubules form dynamic and persistent fibres which survive throughout spindle oscillations. It was suggested that parallel microtubules take turn in maintaining contact with the cell cortex; while a long filament enters a shrinking phase after undergoing catastrophe at the cortex, a neighbouring filament undergoes persistent growth and thereby rescues the fibre. However, whether the dynamics of the different microtubules forming a bundle occur in a coordinated manner or not, and if so, what possible mechanisms might underlie it, are unclear.

In a recent paper, Jemseena and Gopalakrishnan [12] showed that the net polymerisation force of a two-filament bundle consisting of rigid parallel filaments is, in general, sub-additive owing to the “diffusive interaction” between the growing tips of the filaments. Here, the term diffusive interaction refers to the following effect: the growth-rate of (any) one filament is dependent on the location of the tip of the other filament, since both the tips are competing for monomers from the same pool. Hence, the effective single-filament growth rate in a two-filament bundle is dependent on the base separation between the filaments, and is an increasing function of the same. Consistent with this expectation, it was confirmed in numerical simulations that the net polymerization force generated by this two-filament bundle is smaller than twice the force generated by a single filament. This remarkable non-additivity of polymerisation force is also consistent with earlier experimental observations[9] on polymerisation force of a single microtubule, which was found to be much smaller than the maximum (additive) force originating from the 13 protofilaments.

The motivation for the work reported here arises from the above observations, and may be stated clearly as follows. The catastrophe frequency of a microtubule is known to depend strongly on the growth rate (which can be observed through a dependence of the former on the tubulin concentration in solution [13,14]). Therefore, it seems plausible that the catastrophe frequency for a filament which is part of a bundle should also depend on the distance to the neighbouring filament(s). Here, we confirm this effect by performing numerical simulations with a simple one-dimensional stochastic model of microtubule filaments in various bundle configurations, with Brownian diffusion of tubulin monomers explicitly included. Our results suggest that catastrophe events in neighbouring parallel filaments occur in a correlated manner, which could have implications for the dynamics of microtubule filaments in a bundle.

### 1.1 Model Details

For our numerical simulations, we use a one-dimensional model of a microtubule, similar to what has been used in a few earlier studies[15–19]. In this model, a single microtubule is a straight, rigid polymeric filament, which grows by the attachment of (GTP-bound) tubulin at its tip. A free tubulin monomer is assumed to be a point particle which diffuses with diffusion coefficient *D* inside a rectangular box of dimensions *L* × *L* × *H*, one face of which contains the nucleation sites for microtubules. The binding of a tubulin monomer to the tip of a filament is diffusion-limited; after incorporation in a polymer, the monomer is visualised as a solid cylinder of radius *a*. Once a tubulin molecule binds to the tip, the length of the polymer increases by one monomer length. In order to approximately account for the 13-protofilament structure of a microtubule, we assume that a single monomer’s length *ℓ* is 1/13 of the length of a tubulin dimer (8 nm). A polymer with *m* ≥ 1 monomers is assumed to have the shape of a solid cylinder of radius 12 nm and length *mℓ*.

A polymerized GTP-tubulin may undergo spontaneous and irreversible hydrolysis and become GDP-tubulin with rate *r*. This could happen anywhere in the filament. We assume that the free GTP-tubulin is prohibited from binding to a GDP-tubulin; this means that once a filament undergoes catastrophe, further growth of the filament cannot take place. The filament then enters a state of shrinking, accompanied by release of GDP-bound tubulin to the solution (which are not tracked) with rate *k_d_*. The shrinking continues until the filament completely disappears at the nucleation site or until an inner island of GTP-tubulin is exposed, whichever comes earlier. The latter events may be characterised as rescue, but we find that they are rare, and typically a filament which undergoes catastrophe will shrink entirely. In such cases, we ensure that another filament starts growing again at the same location without delay.

At the start of the simulation, our box is populated with *N_m_* monomers, placed at various random locations, which then start diffusing. The equation of motion of a single particle is given by the discrete form of the standard over-damped Langevin equation

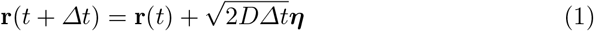

where **r**(*t*) is the three-dimensional position vector of the diffusing particle and ***η*** = (*η_x_, η_y_, η_z_*) is a set of Gaussian random numbers with zero mean and unit variance, representing thermal noise. In our simulations, we used *Δt* = 10^−4^s. The polymer tips act as absorbing surfaces for the diffusing monomers. In order to eliminate any effect due to global depletion of free tubulin due to filament growth, we ensure that for every tubulin molecule adsorbed at the tip of a growing filament, another one is added to the system at a random location on the boundary of the rectangular box. Hence, the number of free monomers remains constant in our simulations.

We carried out simulations of bundles of *N* microtubules, with *N* taking values 2,4 and 5, while *N* = 3 was avoided for reasons of symmetry (since the confining box is rectangular in shape, the three filaments whose bases form an equilateral triangle will not be equidistant from the walls). For *N* = 2 and 4, the filaments are equidistant from each other, with base separation *d*, which is varied. This means that a bundle with *N* = 2 consists of two filaments whose bases are placed symmetrically in the middle of the relevant face of the box, such that the mid-point of the line joining the bases coincides with the geometric centre of the face. Likewise, the configuration *N* = 4 consists of 4 parallel microtubules which nucleate from the four corners of a square of side *d*, such that the centre of the square coincides with the centre of the box. For *N* = 5, we place an additional filament at the centre of the square. To emphasise this difference, the five-filament bundle shall be referred to as “N=4+1”, while the others are designated as *N* = 2 and 4. A schematic figure showing the cross-sections of the various configurations is provided in Fig. 1.

**Fig. 1.**
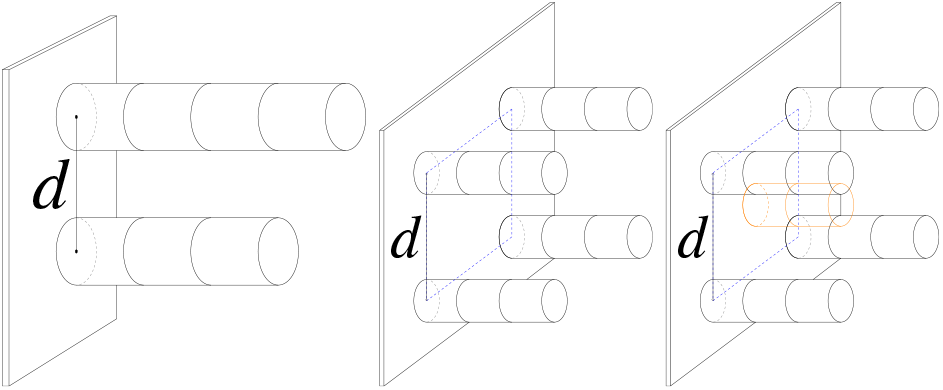
The figure illustrates the geometrical arrangements in *N* = 2, *N* = 4 and *N* = 4+1 configurations of the microtubule bundle.

Table 1 lists the numerical values of all the parameters used in our simulations. Note that a smaller box was used for simulations with more than 2 filaments to reduce computation time.

**Table 1.** List of important parameters used in the numerical simulations.

| Parameter | symbol | numerical value |
| --- | --- | --- |
| Monomer diffusion coefficient | $D$ | $10^5 \text{ nm}^2\text{s}^{-1}$ |
| Simulation box height | $H$ | $1.5\mu\text{m}$ |
| Simulation box side length | $L$ | $3\mu\text{m}$ ($N = 2$)<br>$1.5\mu\text{m}$ ($N > 2$) |
| Spontaneous hydrolysis rate | $r$ | $0.5\text{s}^{-1}$ [19] |
| Detachment rate | $k_d$ | $300\text{s}^{-1}$ [3, 7, 19] |
| Radius of the monomer disc | $a$ | 12 nm |
| Monomer length | $\ell$ | 0.6nm |
| Total number of free monomers | $N_m$ | 2000 |

Fig. 2 shows a typical length versus time trajectory in the two-filament bundle, for base separation *d* = 2nm. Fig. 3 shows the same, when the base separation is larger, with *d* = 70nm. It is interesting to note the obvious difference between the statistics of occurrence of catastrophe events in both the cases. For small base separation (Fig. 2), we observe that between two consecutive catastrophes in any one filament, there are typically multiple catastrophes in the other filament, suggesting that after one catastrophe event has occurred in one filament, the next catastrophe event is more likely to occur in the same filament. When the base separation is increased to 70 nm (see Fig. 3), this correlation is weakened.

**Fig. 2.**
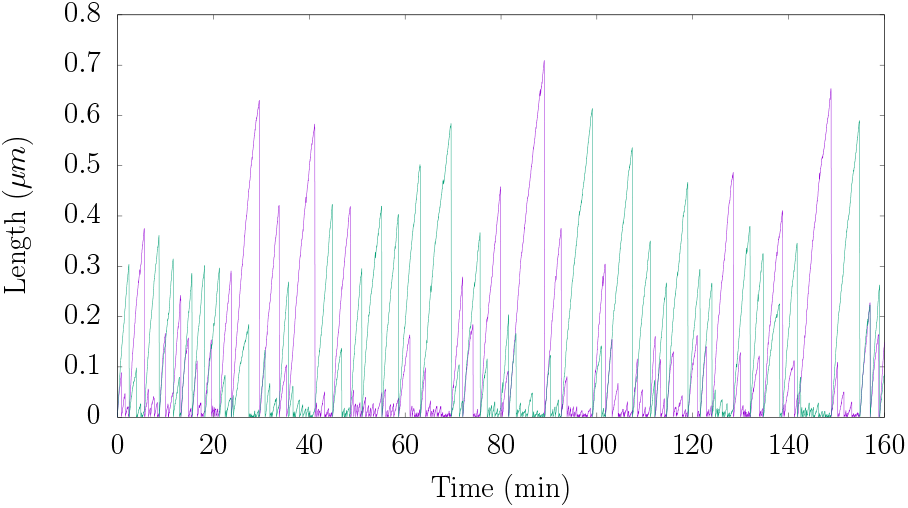
The figure shows length as a function of time for the two microtubules in the *N* = 2 configuration, with the filament bases touching each other, such that base separation *d* = 2nm. Note that multiple catastrophes may occur in one filament between two consecutive catastrophes in the other.

**Fig. 3.**
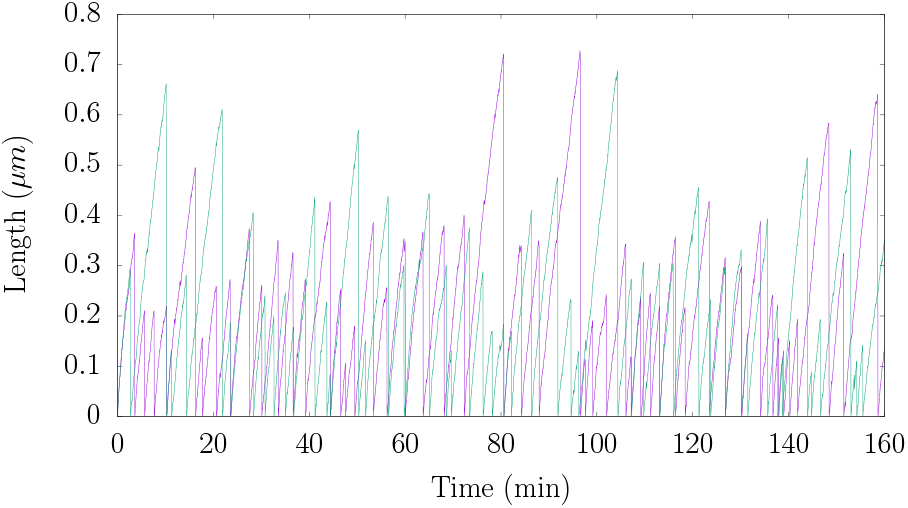
The geometry is similar to that in Fig. 2, but the horizontal separation between the base centres has been increased to *d* = 70nm. Unlike the previous case, the catastrophe events in the two filaments are distributed more or less evenly along time.

We measured the following quantities in our simulations: (a) the mean velocity of growth of a single filament in a bundle (b) the mean time to catastrophe of a single filament (c) a set of correlation coefficients that characterise how likely a catastrophe event in one filament will be followed by a second one in the same filament, or one of the other filaments. The data was collected from 9 different sample systems, each run for 10^4^ seconds.

## 2 Results

### 2.1 Growth velocity and catastrophe frequency

To measure the mean growth velocity of a filament in the growing phase, we identified segments of positive slope in the length versus time plots, as in Fig. 2. The velocity for a given segment with positive slope was found using the chi-squared fitting method. The mean and standard deviation of the velocity was found using the data from different segments. The data is shown in the set of figures Fig. 4 – 6. The plots show that the mean growth velocity is lower when the filaments are closer together. This clearly indicates the presence of diffusive interaction between filament tips, whereby each filament (tip) affects the growth of the other because of overlap of the individual “depletion zones” (regions of space around each tip where the free monomer concentration is lower than the spatial average, due to continuous adsorption). As the filaments are moved farther apart in space, this overlap is reduced, resulting in enhanced growth rate, as the figures show. In all figures, the error bars depict the standard deviation in the measured slope in different growth segments.

**Fig. 4.**
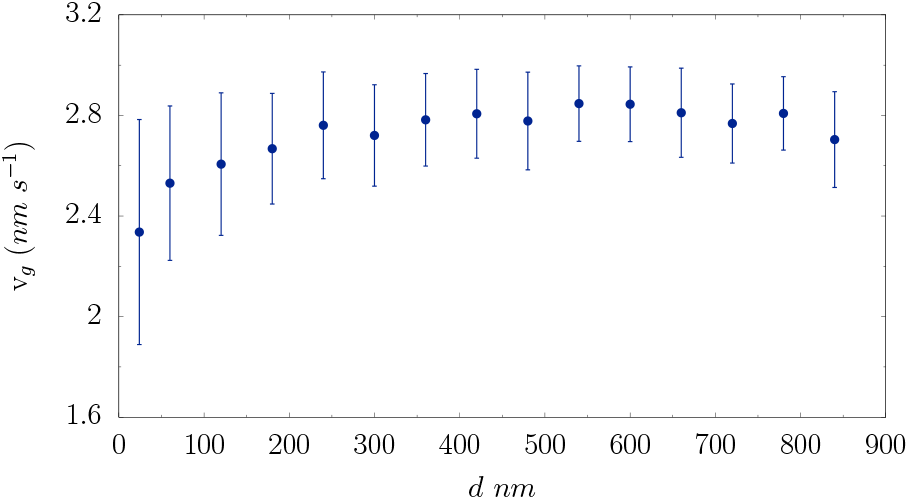
Growth velocity of a filament in a two-filament bundle is plotted as a function of the base separation between the filaments.

**Fig. 5.**
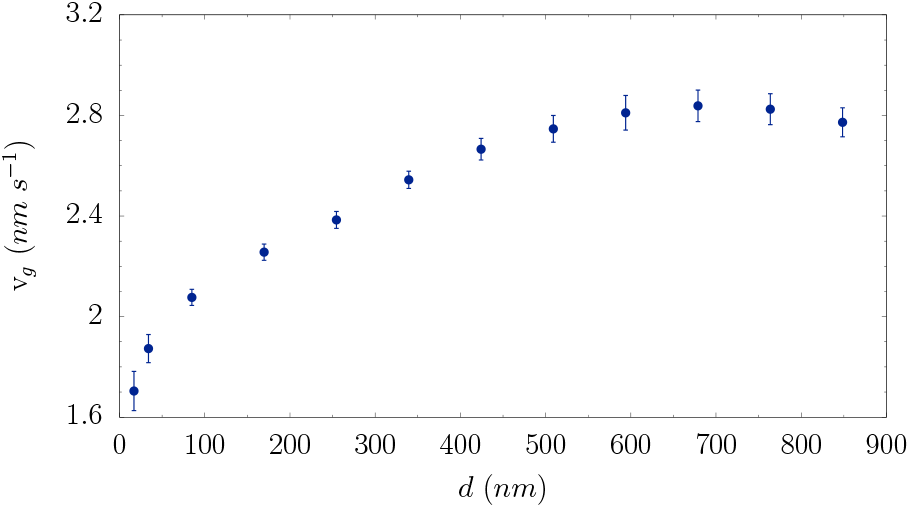
Growth velocity of a filament in a four-filament bundle is plotted as a function of the base separation between the filaments.

**Fig. 6.**
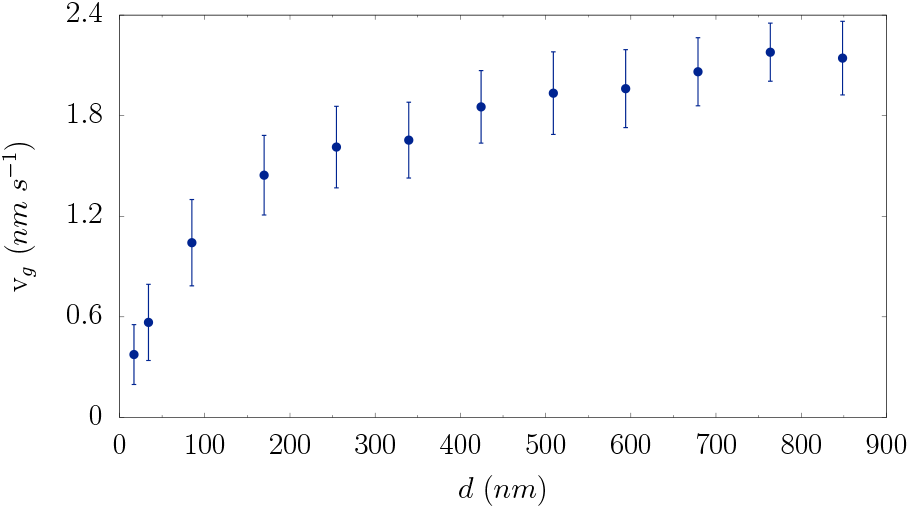
Growth velocity of the centre filament in a bundle in 4+1 configuration is plotted as a function of the base separation between the filaments.

We defined the catastrophe time *T_c_* as the the time interval between the instant a filament starts growing and the instant when it enters a shrinking phase. The inverse of the mean catastrophe time is defined to be the catastrophe frequency *ν_c_*, which is plotted in Fig. 7–9 for various bundle configurations. Typically, a filament that enters the shrinking phase disappears completely, and it is only these events that we considered in computing the catastrophe time. As expected intuitively, the catastrophe time shows an effect similar to growth velocity, hence catastrophe frequency is a decreasing function of the distance between filaments.

**Fig. 7.**
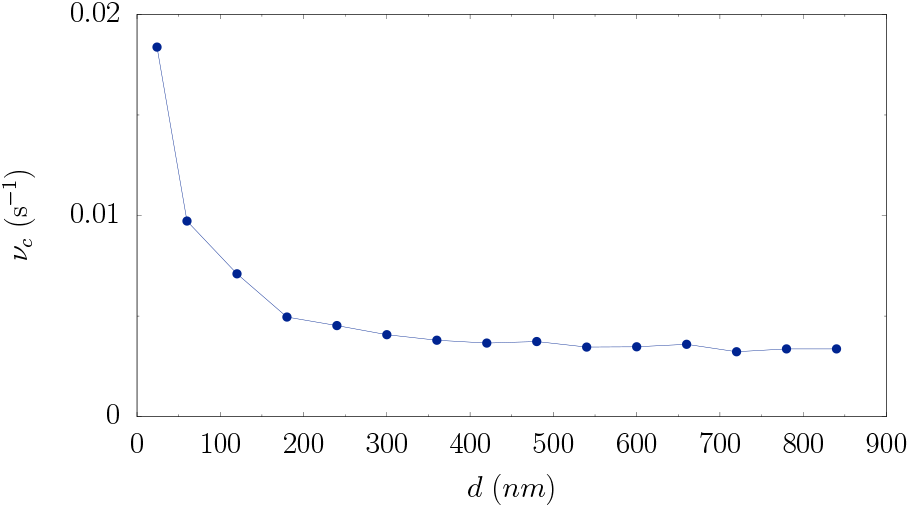
The catastrophe frequency (see definition in text) of a filament in a two-filament bundle are plotted as functions of the base separation between the filaments.

**Fig. 8.**
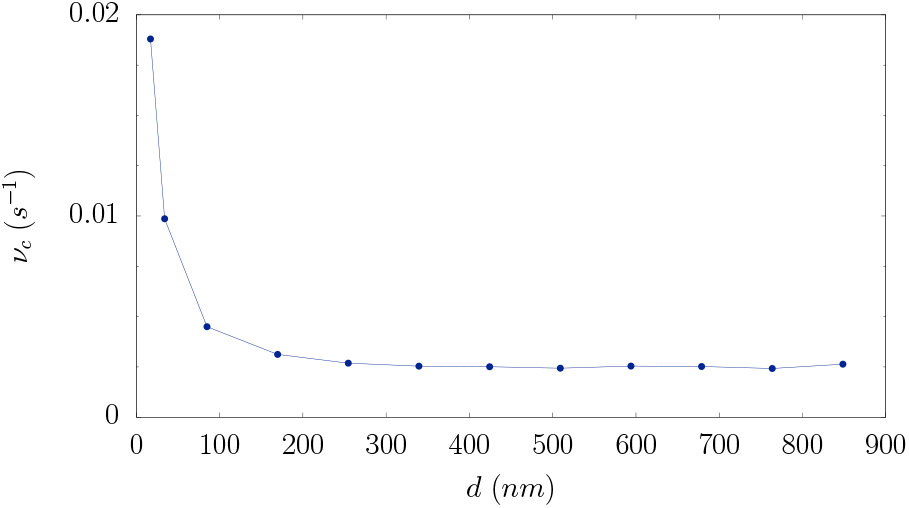
The catastrophe frequency (see definition in text) of a filament in a four-filament bundle are plotted as functions of the base separation between the filaments.

**Fig. 9.**
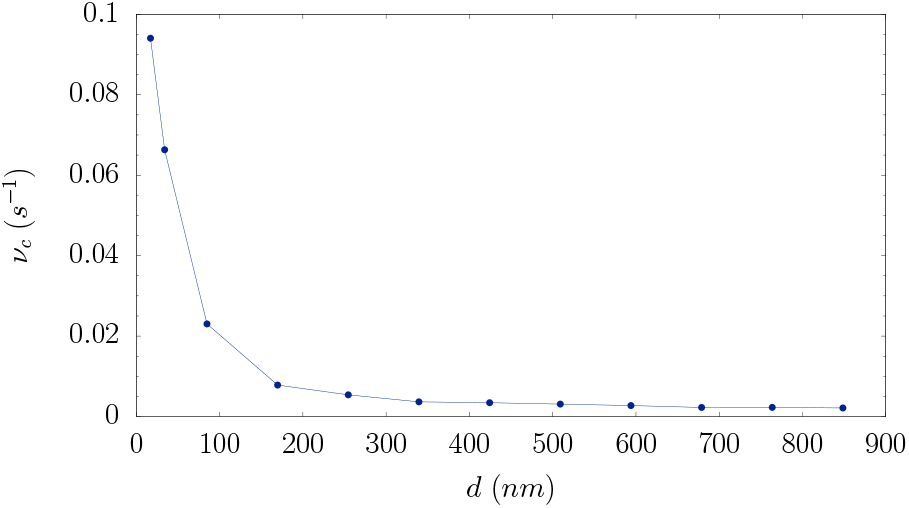
The catastrophe frequency (see definition in text) of the centre filament in a bundle in 4+1 configuration are plotted as functions of the base separation between the filaments.

### 2.2 Correlation in catastrophe events

We defined a set of “splitting probabilities” to characterise the spatio-temporal correlations between catastrophe events occurring in different filaments in a bundle, as follows. In a two-filament bundle, we arbitrarily assigned an identification label *i* = 1 for one of the filaments, while the second filament was designated *i* = 2. The identification of the filaments in the other configurations are indicated in Fig. 10.

**Fig. 10.**
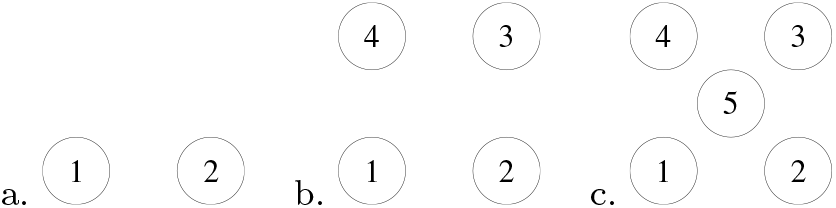
A schematic diagram of the cross-sections of various microtubule bundle configurations studied in this paper, specifying the identification labels for each.

The sequence of catastrophe events observed is now recorded in a binary sequence of the form 12111221…… The information on catastrophe times is disregarded. The number of 12 type intervals (an interval of time, whose origin is a catastrophe event in filament 1, and which ends with a catastrophe event in filament 2, with no catastrophe events in between) is denoted *N*_12_, and similarly we define *N*_11_, *N*_12_ and *N*_22_. The probability that a 1 is succeeded by a 2 is defined as *ϕ*_12_ = *N*_12_/(*N*_12_ + *N*_11_). We may similarly define *ϕ_11_* = 1−*ϕ*_12_, as well as *ϕ*_22_ and *ϕ*_21_ = 1 − *ϕ*_22_. We shall refer to the fractions *ϕ_ij_* as splitting probabilities. Note that since the filaments are identical, we expect *ϕ*_11_ = *ϕ*_22_ and *ϕ*_21_ = *ϕ*_12_. In the absence of any correlation between catastrophe events in different filaments, we expect equal splitting of probability, i.e., *ϕ*_11_ = *ϕ*_12_ = 0.5, hence any deviation in this value indicates that the catastrophe events in different filaments occur in a correlated manner. Specifically, if *ϕ*_11_ > 0.5, we conclude that an event in a certain filament is more likely to be followed by an event in the same filament.

In Fig. 11, we have plotted these probabilities as functions of the base separation *d* between the filaments. It is observed that for small *d, ϕ*_11_ > *ϕ*_12_, and as the distance increases, both quantities approach each other and become equal to 0.5. This indicates the existence of strong correlation between the dynamics of catastrophe events in the two filaments, which is weakened by their increasing separation. For small separations, a certain filament is likely to undergo multiple catastrophes while the other filament undergoes persistent growth. However, since the filaments are identical in every way, symmetry will be maintained and after a while, we expect that the roles will be reversed between the filaments. Our observations in Fig. 2 and Fig. 3 support this picture.

**Fig. 11.**
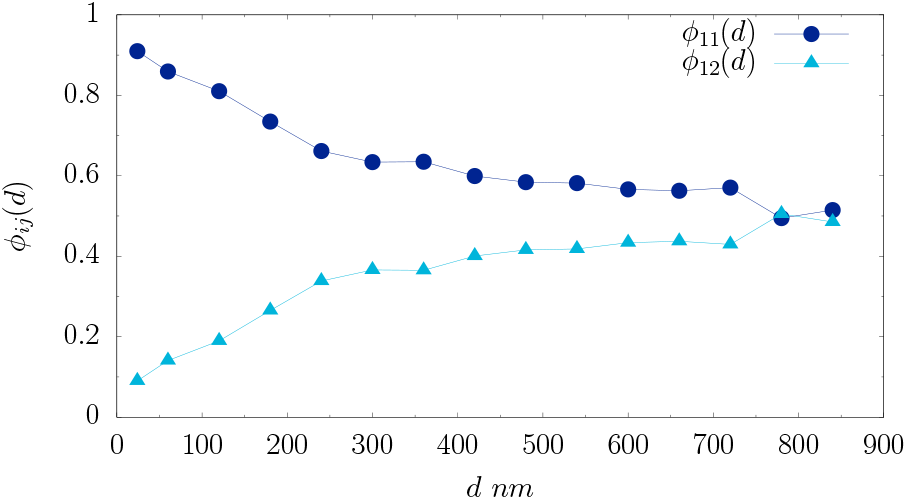
The probabilities *ϕ*_11_ and *ϕ*_12_ in a two-filament bundle are plotted against the filament-filament base separation *d*. For small *d, ϕ*_11_ > *ϕ*_12_, implying a propensity for bunching together of catastrophe events in the same filament, between two catastrophe events in the other filament.

**Fig. 12.**
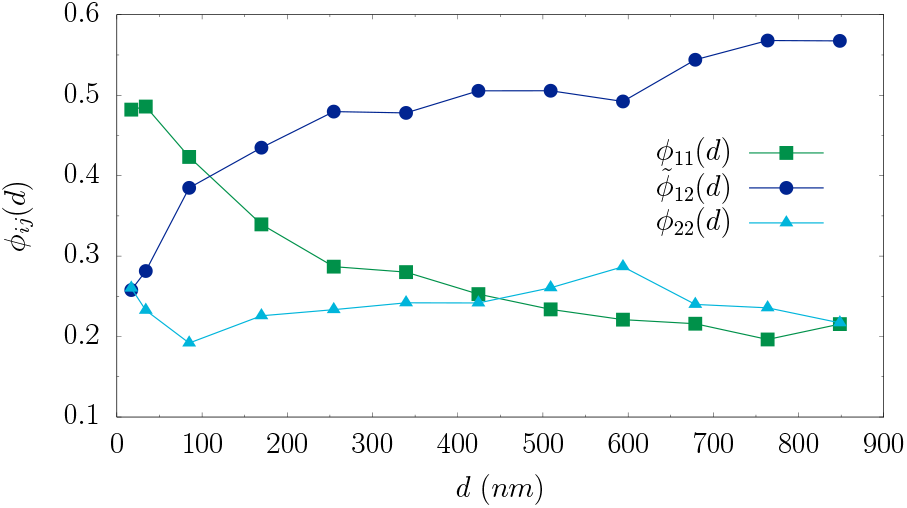
The probabilities *ϕ*_11_, 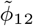 and *ϕ*_13_ in a 4 filament bundle are plotted against the filament-filament base separation *d*. See the text for the definitions of these probabilities.

Why is *ϕ*_11_ > *ϕ*_12_ for small *d* ? Although we do not have a rigorous quantitative explanation, the following argument appears plausible. The two filaments initially grow together, thus creating a tubulin-depleted region around their tips as they grow. A catastrophe in either filament is most probable when the tips are close. Imagine that one filament now undergoes catastrophe and shrinks away. This event releases the other (second) filament from the “influence” of the first one, and the second one will consequently enter a phase of continuous growth. At the same time, the first filament will now shrink to origin and starts to grow again, but suffers from the disadvantage of being trapped in a monomer-poor region. Hence, it suffers multiple catastrophe events before, eventually, it catches up with the first filament and the competition ensues again. Since the filaments are identical in a statistical sense, neither of them has a long-term advantage over the other and reversal of roles happens over a period of time.

The identification of the filaments in the *N* = 4 and *N* = 4+1 configurations is shown in the schematic diagram Fig. 1. In all the cases, we define the probabilities using the formula

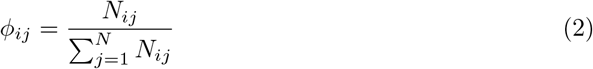

The probabilities are normalised as 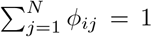 for all *i*. Since filaments 2 and 4 are identical nearest neighbours, when 1 is used as the filament of reference, it is also convenient to define

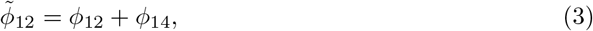

which gives the probability that a catastrophe event in filament 1 is followed by a catastrophe event in either one of the nearest neighbour filaments..

In the *N* = 4 bundle, we observe that *ϕ*_11_ and 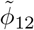 have similar behaviour, as functions of *d*, as the corresponding quantities in the *N* = 2 bundle, while *ϕ*_02_ is more or less independent of *d*. In this case, though, since two filaments (1 and 3) contribute to 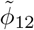 (nearly equally), a crossing between *ϕ*_11_ and 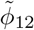 occurs between *d* = 100nm and *d* = 200nm, unlike *N* = 2. There is very little correlation observed between events in 1 and 3, presumably because of the larger separation 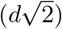.

For *N* = 4 + 1, with the centre filament taken as the reference for reasons of symmetry, the observations are similar to the *N* = 2 and *N* = 4 configurations, as shown in Fig. 13. Here, we show *ϕ*_55_ (the probability that a catastrophe event in the centre filament will be followed by one in the same filament) in comparison with

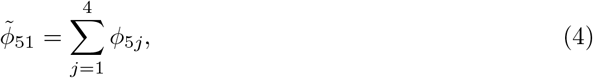

which gives the probability that an event in 5 is followed by an event in one of the (equidistant) corner filaments. For small *d*, 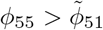, but as *d* is increased, the statistical multiplicative factor of 5 helps the latter overtake the former, and we again encounter a crossing of the probability curves, similar to *N* = 4.

**Fig. 13.**
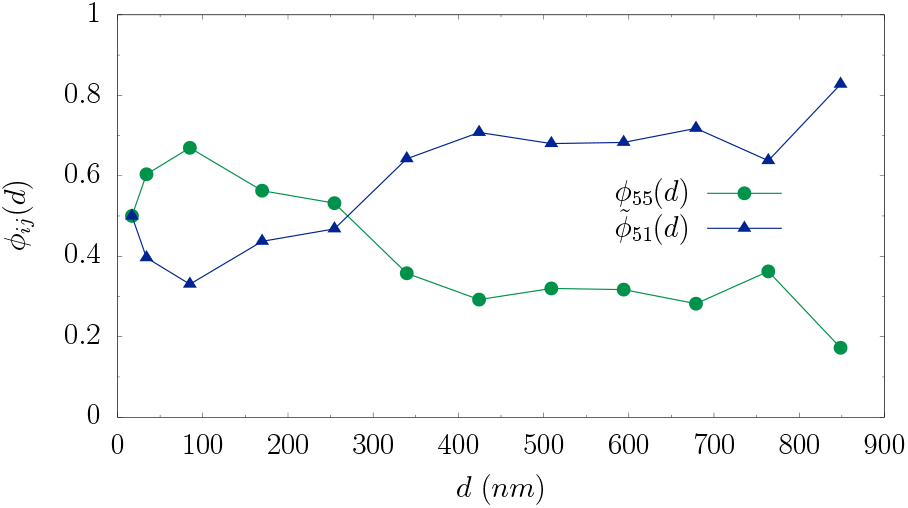
The probabilities *ϕ*_55_ and 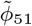 in 4+1 filament bundle are plotted against the filament-filament base separation *d*. See the text for definitions of these probabilities.

## 3 Discussion and conclusions

In this paper, we have investigated the spatio-temporal correlations between catastrophe events occurring in different microtubules that form a bundle, using a onedimensional computational model. In a recent theoretical study, it had been shown that the net polymerisation force produced a bundle of parallel filaments scales sub-linearly with the number of filaments, owing to the presence of diffusive interaction between the filament tips[12]. The diffusive interaction refers to a competition effect between absorbing sinks which irreversibly capture diffusing particles from a common pool, and increases in significance as the sinks come closer together[20]. In the present work, we show that diffusive interaction leads to an an enhancement of catastrophe frequency for individual filaments in the bundle. While this result is along expected lines, a more dramatic manifestation of diffusive interaction appears in the correlation between catastrophe events. Our results demonstrate clearly that when the filaments are closely bundled together, a catastrophe event in one filament is more likely to be followed by another in the same filament. These self-catastrophe events show a propensity for bunching together: two consecutive catastrophe events in any one filament separated by a long interval will likely have multiple catastrophes in the other filament sandwiched between them.

Our results are likely to be significant in understanding the dynamics of microtubule bundles. Microtubule bundles growing against a forced barrier have been observed to undergo collective catastrophes[9] where the individual filaments switch to a state of shrinking in an apparently coordinated manner. We believe that this is likely a consequence of diffusive interaction between the growing tips of different filaments. In the presence of the barrier, the tips are forced to stay close to each other, which enhances the negative effect they exert on each other’s growth and thereby increases the likelihood of catastrophes. In addition, after one filament undergoes catastrophe, the contact time of the rest of the filaments with the barrier would increase on average, and this would lead to more catastrophes. If this argument were true, we should observe the opposite effect in a free bundle; here, the occurrence of a catastrophe event in one filament should have a negative effect on catastrophe in a neighbouring filament by promoting its growth. On an average, this mechanism is beneficial to the growth of a filament bundle by ensuring that while some of the filaments are undergoing catastrophes one after another, the rest of the filaments persist in a state of growth. This is similar to experimental observations on the dynamics of astral microtubules in *C. elegans* embryos[11], where persistent fibres have been shown to be a crucial factor in spindle oscillations. The observed “anti-correlation” between growth of neighbouring microtubules in our simulations might also be relevant for search and capture processes[21–23], where the dynamic instability of microtubules plays a central role.

To conclude, in the present study, we have uncovered a novel mode of interaction between parallel microtubules in a bundle, which leads to correlated growth and catastrophe events. A number of preliminary results have been obtained via Brownian dynamics-based numerical simulations. The findings presented here provide solid evidence for the existence of this interaction and its significance. In future, we plan to expand this study to investigate the effects of this interaction on collective force generation of a parallel microtubule bundle. Experimental validation of these results is highly desirable, as would be a mathematical theory.

## Acknowledgements

M.D acknowledges the P.G. Senapathy Centre for Computing Resources, IIT Madras for time in their Virgo cluster. M.G would like to thank Chaitanya Athale for fruitful discussions.

